# An ancient transcription activator is required for the multiciliogenesis program

**DOI:** 10.1101/2023.12.16.571971

**Authors:** Haochen Jiang, Mengting Hua, Mei Ding, Jiaqi Hu, Shiqiong Zheng, Hanchen Wang, Weihua Wang, Cheng Xu, Junqiao Xing, Hongni Liu, Xue Zhao, Zhangfeng Hu

**Affiliations:** Institute of Biomedical Sciences, School of Medicine, Jianghan University, Wuhan, Hubei 430056, China; Hubei Engineering Research Center for Protection and Utilization of Special Biological Resources in the Hanjiang River Basin, School of Life Sciences, Jianghan University, Wuhan, 430056 Hubei, China

**Author notes:** Correspondence (H.J.), (Z.H.). These authors contributed equally (Haochen Jiang, Mengting Hua, Mei Ding, Jiaqi Hu).

## Abstract

Multiciliogenesis is an evolutionarily conserved process required a unique transcriptional program. The mechanisms governing multiciliogenesis are incompletely understood. Here we show that an ancient transcription activator, Edf1, is essential for the multiciliogenesis program. Mice lacking Edf1 exhibit postnatal hydrocephalus and delayed multiciliated cells (MCCs) differentiation. Functional studies reveal that MCCs in Edf1^-/-^ mice have defects in tissue-level polarity and reduced motility. These defects result in abnormal cerebrospinal fluid (CSF) dynamics in vivo, potentially contributing to the development of hydrocephalus. Intriguingly, we found that the expression of pivotal ciliary transcription factors was decreased in Edf1^-/-^ mice. Collectively, our data suggest that Edf1 modulates the multiciliogenesis by regulating the transcription of important ciliary transcription factors required for ciliary gene expression.

## Introduction

Cilia and flagella are ancient organelles present in almost all groups of eukaryotes. Although many eukaryotic cells exhibit only one cilium, specialized cells possess more than two cilia, collectively known as multiciliated cells (MCCs). In unicellular organisms, multi-cilia organelles are essential for feeding, sensation, and locomotion. In vertebrates, MCCs line the luminal surface of diverse tissues (Marinković, Berger, & Jékely, 2020). Dysfunction of MCCs is implicated in a broader spectrum of human diseases (Defosset et al., 2021; Lovera & Lüders, 2021; Reiter & Leroux, 2017). Despite MCCs playing pivotal roles in health, they remain relatively understudied.

Congenital hydrocephalus is a neurological disorder characterized by the accumulation of excessive CSF that forces the ventricles to expand, leading to a series of brain function problems with high mortality (Casey et al., 1997; Miyan, Nabiyouni, & Zendah, 2003; Vogel et al., 2012). The classical pathogenic mechanism leading to hydrocephalus is the damage to cerebrospinal fluid circulation caused by ciliary-driven flow defects or imbalanced absorption (Spassky & Meunier, 2017). Defects in ependymal cell differentiation or ciliary movement may lead to excessive accumulation of cerebrospinal fluid and induce hydrocephalus (Del Bigio, 2010; Garcia-Bonilla, McAllister, & Limbrick, 2021). Therefore, identifying the pathogenic genes related to congenital hydrocephalus induced by defects in multiciliated ependymal cells (MCECs) development becomes important to prevent genetic defects in patients.

How the transcriptional control of multiciliogenesis evolved is an outstanding question in MCC biology. Transcriptional regulation of ciliary genes is an important way to achieve multiciliogenesis control in both unicellular and multicellular organisms (Choksi, Lauter, Swoboda, & Roy, 2014; Lewis & Stracker, 2021). Recently, several transcription regulators were identified as central players required for multiciliogenesis in multicellular organisms. However, multiple key ciliary transcription regulators, including Gemc1 (Gmnc), Mcidas (Multicilin), Foxj1 and RFXs, appear to be absent in unicellular MCCs (Defosset et al., 2021; Zhou et al., 2015).

Edf1 (Endothelial differentiation-related factor 1) proteins are evolutionarily conserved across diverse organisms including both unicellular and multicellular organisms. Accordingly, Edf1 is a transcriptional co-factor that participates in the differentiation process of various cells (Busk et al., 2003; Leidi, Mariotti, & Maier, 2009; Mariotti, De Benedictis, Avon, & Maier, 2000; K. Takemaru, Harashima, Ueda, & Hirose, 1998; Ki Takemaru, Li, Ueda, & Hirose, 1997). However, whether Edf1 is involved in central nervous system development, especially the development of MCECs in vivo is still unknown. In the present study, we found that Edf1 protein expression is present in RGCs and MCECs. Edf1 knockout mice showed runted and postnatal hydrocephalus. Edf1 knockout impairs MCEC development. In addition, Edf1 knockout MCECs exhibit shorter cilia length, and dysfunction of ciliary movement, resulting in abnormal CSF mediated by ependymal cells. Mechanistically, Edf1 positively regulates transcription networks in the early stages of MCEC differentiation. Our results reveal the important role of Edf1 in the early transcriptional regulation process of MCEC development and suggest that Edf1 might be a candidate gene for human congenital hydrocephalus disorder.

## Results

### *Edf1* gene deletion causes postnatal hydrocephalus

Edf1 is highly conserved across diverse organisms including both unicellular and multicellular organisms (Figure 1-figure supplement 1A and B). To investigate the function of Edf1 protein, we generated *Edf1* knockout mice using the CRISPR-Cas9 system. Deletion of Edf1 was confirmed by immunoblotting and RT-qPCR of several tissues (Figure 1-figure supplement 1C-E). Heterozygous *Edf1*^+/-^ mice appeared normal (Figure 1-figure supplement 1F). Interbreeding *Edf1*^+/-^ mice produced offspring that appeared normal at birth. However, only 38% of *Edf1*^-/-^ mice survived after the first two weeks postnatal compared to the wild type, then survived *Edf1*^-/-^ mice were viable for at least 8 months (Figure 1-figure supplement 1G and Figure 1-figure supplement Table 1). At postnatal day 3 (P3), the body size and weight of *Edf1*^-/-^ mice were lower than those of wild type littermates (Figure 1A-C). At about P15, *Edf1*^-/-^ mice exhibited cranial protrusions. Histological analysis of *Edf1*^-/-^ mouse brains using sequential hematoxylin and eosin (H&E) stained coronal sections revealed that the lateral ventricle and third ventricle (LV and 3V, respectively) were enlarged in *Edf1*^-/-^ brains, whereas no ventriculomegaly was detected in the fourth ventricle (4V) (Figure 1E). Expansion of the ventricles was first observed in *Edf1*^-/-^ brains at P3 (Figure 1-figure supplement 1H) and maximal extension was reached at P16-P20, suggesting that *Edf1*^-/-^ mice develop hydrocephalus postnatally. From P3 to P128, we examined the hydrocephalus of *Edf1*^-/-^ brains and found that only one mouse did not develop hydrocephalus and had normal body size (Figure 1D). These results established that Edf1 was required for normal growth and has a strong effect on brain development.

**Figure 1.**
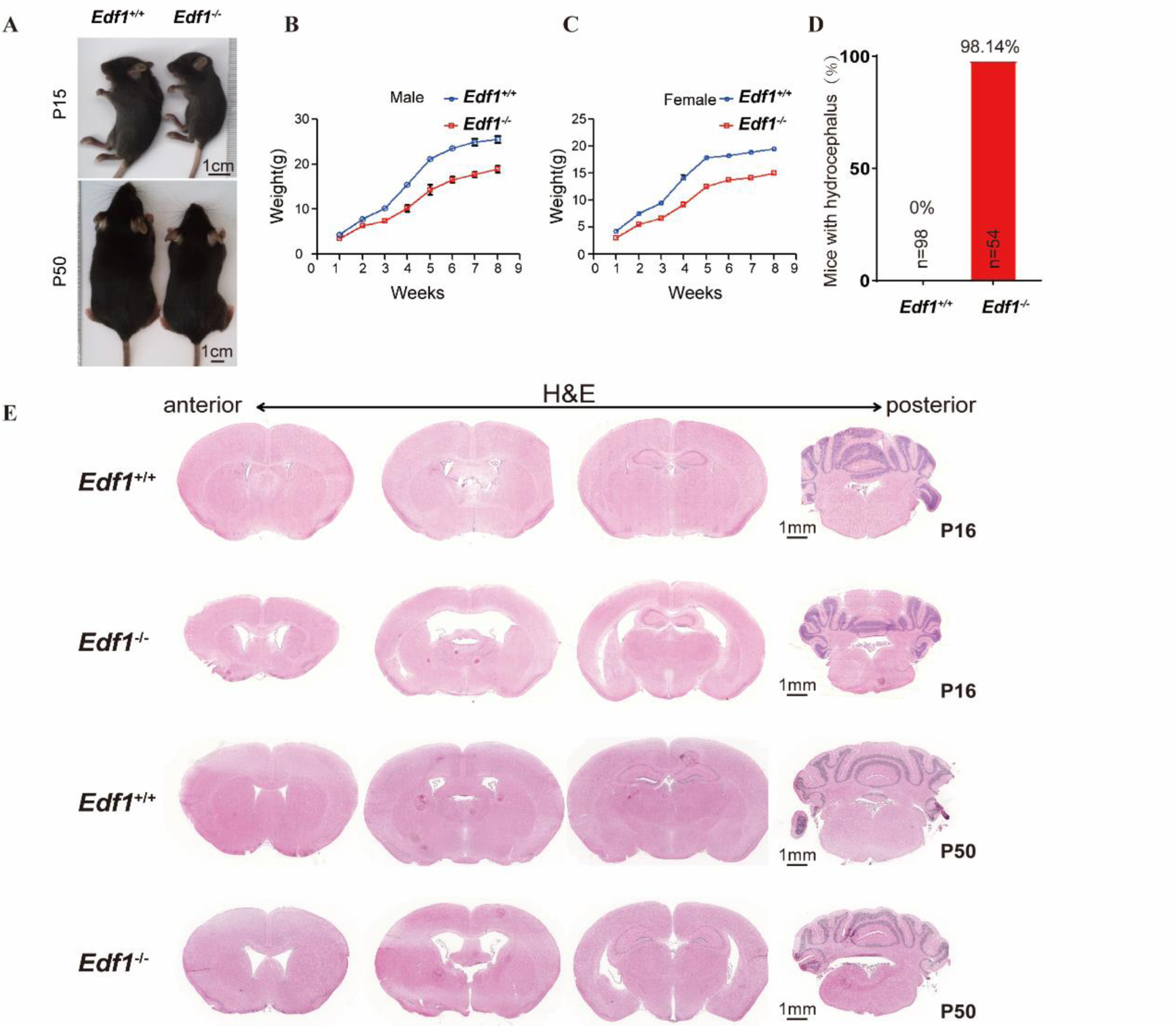
E*d*f1 gene deletion causes postnatal hydrocephalus. (A) Littermate males of the wild type or *Edf1* knockout at indicated ages postpartum. Scale bar: 1 cm. Additional examples are provided in Figure 1-supplement Figure 1F. (B-C) Weights of mice of wild type or *Edf1* knockout from 1 to 8 weeks. (B) male 5 mice for each genotype. (C) female 8 mice for each genotype. Error bars indicate ± SEM. (D) Quantification of hydrocephalus incidence in wild type and *Edf1*^−/−^ mice. Mice number as indicated. (E) H&E stained coronal sections of the wild type and *Edf1*^−/−^ mice brains at indicated ages postpartum. Scale bar: 1mm

### Anatomical analysis of *Edf1*^-/-^ brain

Cortical hypoplasia and stenosis in the Sylvian aqueduct are frequently associated with congenital hydrocephalus (Casey et al., 1997). Before and after the onset of hydrocephalus, the Sylvian aqueduct was not significantly different between *Edf1*^-/-^ and wild-type controls (Figure 2A). The subcommissural organ (SCO) is a secretory gland positioned immediately anterior to the Sylvian aqueduct underneath the posterior commissure. SCO agenesis results in hydrocephalus (Huh, Todd, & Picketts, 2009). In *Edf1*^-/-^ mice, the SCO had a similar size compared to that in controls (Figure 2B. The morphology of the choroid plexus and the expression of choroid plexus proteins AQP1 and β-catenin was normal in the choroid plexus of control and *Edf1*^-/-^ mice (Figure 2D). Interestingly, although *Edf1*^-/-^ mice show small body size after birth, there is no significant difference in cortical thickness during development (Figure 2C). Furthermore, the localization of β-catenin at cellular membranes of LV was not structurally altered, indicating that cell-to-cell adhesions and tight junctions were not affected by Edf1 deletion (Halbleib & Nelson, 2006) (Figure 2-figure supplement 1). According to the results, we suggest *Edf1* knockout in mice leads to the formation of hydrocephalus in the brain postnatally and is not caused by tissue structural defects.

**Figure 2.**
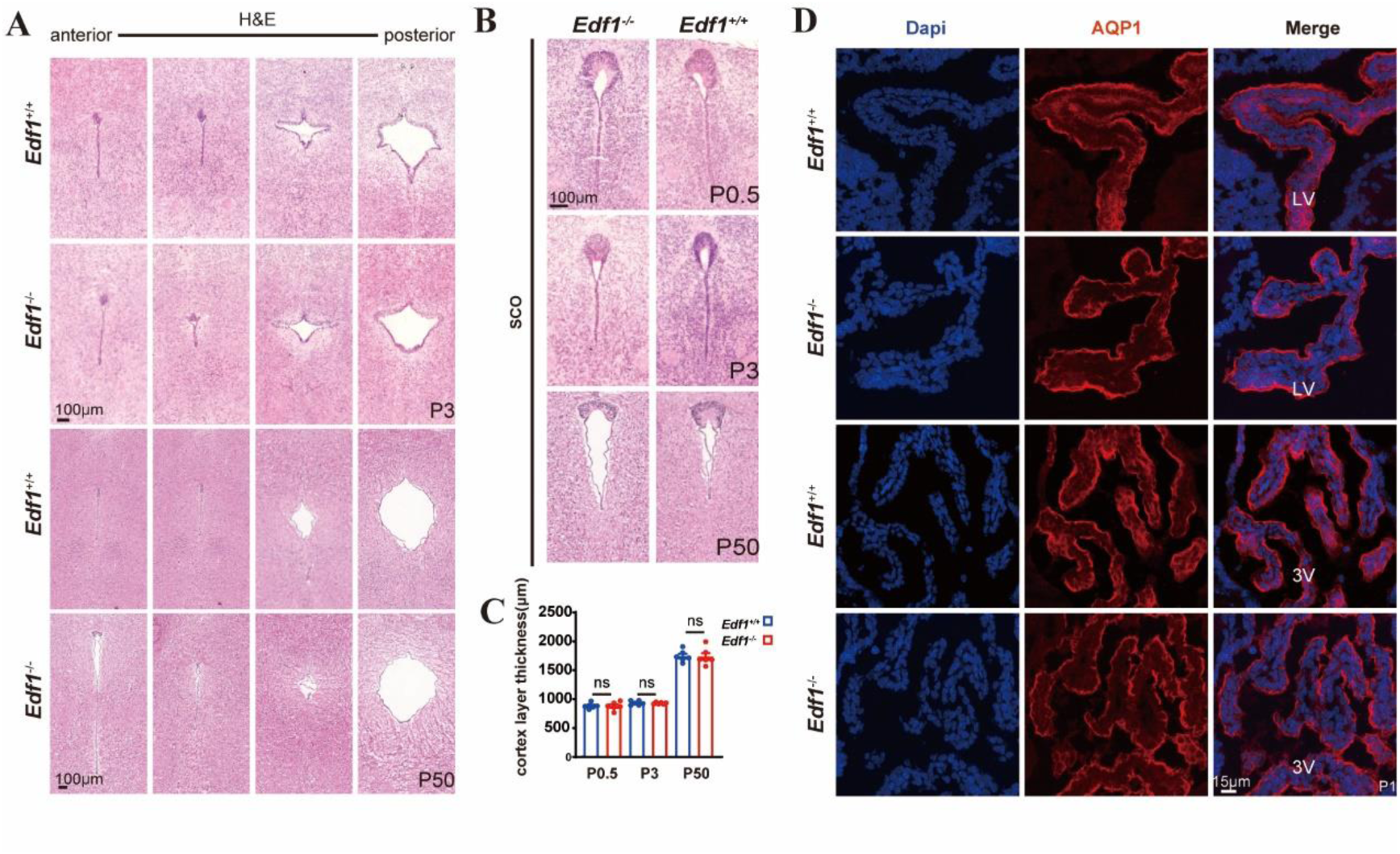
Anatomical analysis of the *Edf1* knockout brain. (A) Coronal sections in the Sylvian aqueduct of wild type and *Edf1*^−/−^ mice at indicated ages. 100 μm intervals from anterior (left) to posterior (right) for P3.5, and 200 μm for P50. Stenosis was not observed in any section of each genotype at indicated ages. Scale bar: 100 μm. (B) Similar morphology of SCO structure of wild type and *Edf1*^−/−^ mice at indicated ages. Scale bar: 100 μm. (C) Cortex layer thickness of wild type and *Edf1*^−/−^ mice showed no significant differences at indicated ages. n=6 for each genotype at indicated ages. Error bars indicate ± SEM. All *P*-values were calculated using an unpaired, two-tailed Student’s t-test. ns., not significant. (D) Immunohistochemical analysis of the choroid plexus of LV and 3V. The wild type and *Edf1*^−/−^ choroid plexus were stained with anti-AQP1 (red) and Dapi (blue). Scale bar: 15μm.

### Edf1 expression in embryonic and postnatal brain

Edf1 is highly expressed in brain tissue (Figure 1-figure supplement 1D). To further explore how Edf1 deficiency affects brain tissue development and leads to hydrocephalus, we performed Edf1 immunofluorescence staining on embryonic and postnatal brains. In the embryonic central nervous system, Edf1 is mainly expressed around the ventricles and the outer edge of the cerebral cortex. In postnatal mouse brains, Edf1 protein is more widely distributed and is localized in the cerebral cortex, hippocampus, striatum, SCO, choroid plexus, and ependymal cells of the ventricles and aqueducts (Figure 3A).

**Figure 3.**
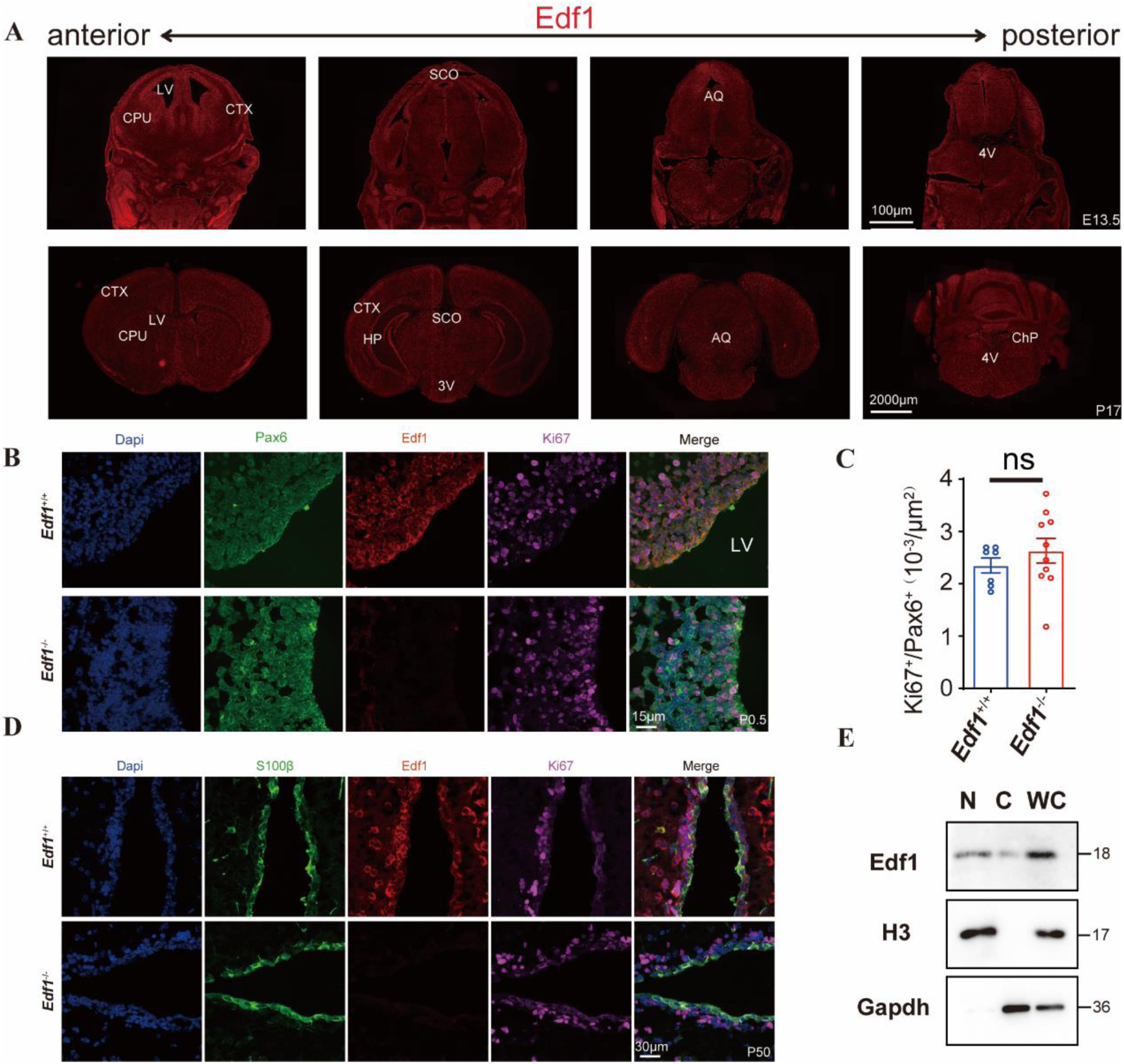
Edf1 expression in embryonic and postnatal brain. (A) Immunofluorescence labeling of Edf1 was performed on coronal sections of E13.5 (top panel, Scale bar: 100 μm) and P17 brain (bottom panel, Scale bar: 2000 μm). ChP choroid plexus, CTX cerebral cortex, CPu Caudate Putamen, LV lateral ventricle, 3V third ventricle, 4V fourth ventricles Hip hippocampal formation. (B-C) Coronal brain sections from wild type and *Edf1*^−/−^ mice at P0.5 were stained for anti-PAX6(green), anti-Edf1(red), anti-Ki67 (magenta), and Dapi (blue) with associated cell counts. For cell counts,7 sections from 3 mice of wild type and 10 sections from 5 *Edf1*^−/−^ mice were counted. Scale bar: 15 μm Error bars indicate ± SEM. All *P*-values were calculated using an unpaired, two-tailed Student’s t-test. ns., not significant. (D) Coronal brain sections from wild type and *Edf1*^−/−^ mice at P50 were stained for anti-S100β(green), anti-Edf1(red), anti-Ki67 (magenta), and Dapi (blue). Scale bar: 30 μm. (E) Western blot analysis showing the subcellular localization of endogenous Edf1 in RGCs. H3 is used as a nuclear protein marker, and Gapdh is used as a cytoplasmic protein marker.

Furthermore, immunofluorescence co-localization showed that during the perinatal period, Edf1 had a significant co-localization relationship with Pax6, a radial glial cell marker, in the subventricular zone (SVZ) (Figure 3B). Given that RGCs become committed and differentiate into various neuronal lineages, we subsequently confirmed the Edf1 expression lineage in postnatal brain tissue. On the surface of the ventricles, Edf1 had a co-localization relationship with S100β, which is expressed in ependymal cells (Figure 3D). In addition, in the cortex and hippocampus, Edf1 co-localizes with NeuN, a mature neuronal marker (Figure 3-figure supplement 1A and B). Early abnormal proliferation or development of neurons can lead to structural defects in brain tissue, which in turn can cause hydrocephalus (Duy et al., 2022; Ito et al., 2021; Jin et al., 2020). Immunofluorescence staining using the proliferation marker Ki67 can reveal that neural proliferation in the SVZ region is not significantly different between *Edf1*^-/-^ mice and wild-type mice (Figure 3C). Cell proliferation ability detected by EDU in vitro isolation of neural stem cells is unaffected in *Edf1*^-/-^ mice (Figure 3-figure supplement 1D and E). By counting the number of NeuN in the cortex and hippocampus of wild-type and *Edf1*^-/-^ mice, it is shown that mature neuronal development is not impaired in *Edf1*^-/-^ mice (Figure 3-figure supplement 1C). Based on these results, it can be concluded that hydrocephalus induced by *Edf1* knockout is not due to impaired neurogenesis. Previous studies have shown that Edf1 acts as a transcriptional co-factor involved in various physiological metabolic processes (Ballabio, Mariotti, De Benedictis, & Maier, 2004; Leidi et al., 2009; Mariotti et al., 2000). By performing nuclear-cytoplasmic separation experiments, we can see that Edf1 is expressed in both the nucleus and cytoplasm of RGCs in vitro (Figure 3E).

### The *Edf1* knockout results in an abnormal development of multiciliary ependymal cells

Defects in the differentiation and ciliogenesis of ependymal cells can lead to hydrocephalus (Ji et al., 2022). Given the onset of hydrocephalus in *Edf1*^-/-^ mice is consistent with the time of RGC-to-mature ependymal differentiation (after P2), we suspect that hydrocephalus induced by *Edf1* knockout may be due to abnormal development of ependymal multiciliated cells (Spassky et al., 2005). We selected the P3 period developing brain because it is in the early stage of *Edf1* knockout-induced hydrocephalus and also the early stage of MCEC development in ependymal cells. This is conducive to our intuitive discovery of any abnormal cilia development and to avoid any subsequent effects caused by hydrocephalus.

Scanning electron microscopy (SEM) was used to observe the development of MCECs in the lateral ventricle ependyma of *Edf1^-/-^* mice at P3. Ependymal cilia begin to develop along the lateral wall of the ventricle in the caudo-ventral to rostro-dorsal direction from birth (Delgehyr et al., 2015). To avoid observation errors caused by different observation positions, we selected three positions along the anterior-posterior direction of the lateral ventricle. In P3 wild-type controls, most of the caudal ventricular walls (Point2) were covered with cilia tufts, while the density of cilia tufts in both lateral positions was lower (point 1,3). However, in *Edf1^-/-^* mice, there was a significant decrease in cilia tufts density in comparable ventricular regions at P3, especially in rostral areas where there were almost no cilia (Figure 4A). Using whole-mount staining for β-catenin and γ-tubulin, we observed pinwheel organization with large apical surfaces and multiple basal bodies in the caudal ventricular walls of wild type control. In contrast, in *Edf1^-/-^* mice, there were still many undifferentiated ependymal cells in the same area (Figure 4B).

**Figure 4.**
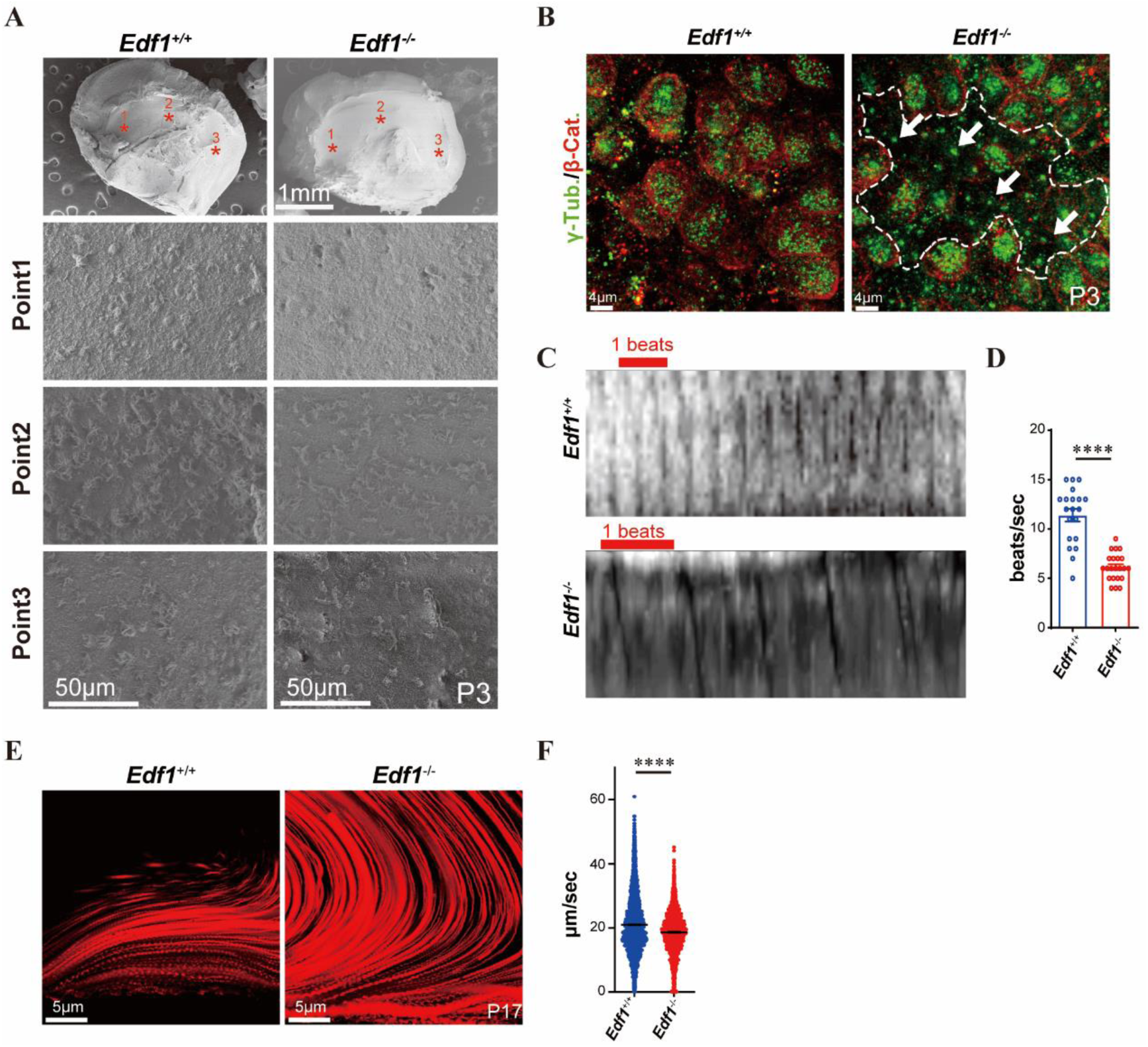
The *Edf1* knockout results in an abnormal development of multiciliary ependymal cells. (A) SEM was used to visualize the ultrastructure of the ependymal cilia in wild type and *Edf1*^−/−^ lateral ventricles at P3. The top panel indicates the areas of selection at P3. Point 1 represents the anterior part of the lateral ventricle walls. Representative images were taken with 2000x. Scale bar: 50 μm. (B) Wholemount immunostaining for β-catenin(red) and γ-tubulin (green) of wild type and *Edf1*^−/−^ lateral ventricles at P3. The white dotted line area of *Edf1*^−/−^ lateral ventricles represents undifferentiated MCECs. White arrows indicate RGCs which have a small apical surface with a single basal body. Scale bar: 4 μm. (C) Sequential images of ciliary beating in wild type and *Edf1*^−/−^ mice at P5 for 1 sec. The red solid line indicates a single swing. (D) Ciliary beating frequency in wild type (n = 20 from 10 whole-mount preparations, 5 mice) and *Edf1*^−/−^ mice (n = 23 from 13 whole-mount preparations, 7 mice). Data shown are means ± SEM. Each point on the graph is the average rate of ciliary beating in a given field of individual MCECs. (E) Migration patterns of the fluorescent beads placed in the LV whole-mount preparations of wild type and *Edf1*^−/−^ mice. Red represents the trajectory of the fluorescent beads within 30 sec. (F) The speed of fluorescent beads was slower in *Edf1*^−/−^ mice than wild type mice (n = 3469 beads from 8 wild type mice, n = 2557 beads from 8 *Edf1*^−/−^ mice). Each point on the graph is the speed of individual fluorescent beads. Error bars indicate ± SEM. All *P*-values were calculated using an unpaired, two-tailed Student’s t-test, \*\*\*\**P* < 0.0001.

The swinging of cilia on the apical surface of ependymal cells to produce unidirectional fluid flow is crucial for generating ependymal flow (Ji et al., 2022). Ciliary beating frequency was directly observed using high-frequency cameras in fresh coronal brain slices containing the lateral ventricles. The ciliary beating frequency in *Edf1^-/-^*mice was significantly lower than that in the control group (Figure 4C and D, Figure 4-figure supplement movie 1 and 2). These results indicate that *Edf1* knockout leads to abnormal differentiation of MCECs in mice with a decrease in ciliary beating frequency, which may be the cause of early-onset hydrocephalus. It is worth noting that the ventricular dilation in *Edf1^-/-^* mice reaches its maximum at P16-P20. Through whole-mount staining, it was found that at this stage, the density of MCECs in *Edf1^-/-^* mice had reached the level of wild-type controls (Figure 4-figure supplement 1A).

However, by measuring the alignment of basal body rotation known as tissue-level polarity, it was found that coalignment between ependymal cells in *Edf1^-/-^* mice was significantly disrupted (Figure 4-figure supplement 1B and C). Basal bodies (BB) patch displacement used to characterize ciliary beating amplitude (Mitchell et al., 2009; Vladar, Bayly, Sangoram, Scott, & Axelrod, 2012) was also significantly reduced in *Edf1^-/-^*mice (Figure 4-figure supplement 1D). Furthermore, in experiments using fluorescent beads to detect local cerebrospinal fluid flow rate in the lateral ventricle, we found that the flow direction in *Edf1^-/-^* mice was inconsistent and that local flow velocity was also reduced compared to wild-type controls (Figure 4E and F, Figure 4-figure supplement movie 3 and 4). In summary, these results suggest that Edf1 is involved in the development and physiological function of MCECs in vivo.

### Edf1 regulates the ciliogenic program of ependymal cells via regulating ciliary transcriptional networks

To explore the mechanism of Edf1 regulating the development of MCECs, we first wanted to know whether the ependymal developmental defects caused by *Edf1* knockout were due to the loss of endogenous Edf1 in cells. We isolated RGCs from *Edf1^-/-^* mice or wild-type P0 mice and induced them to differentiate into multiciliated ependymal cells in vitro. By immunofluorescence staining of acetylated-tubulin to label cilia, we counted the proportion of MCECs and cilia length. We found the proportion of MCECs formed by *Edf1* knockout was significantly lower than that of wild-type, and the cilia length was slightly reduced (Figure 5A-C). Foxj1 and Rfx2, as key transcription factors for multiciliogenesis (Lemeille et al., 2020; Stubbs, Oishi, Izpisúa Belmonte, & Kintner, 2008; Vij et al., 2012; Yu, Ng, Habacher, & Roy, 2008), increased significantly in protein content with the formation of MCECs during the induction process of wild type, but decreased significantly in *Edf1* knockout (Figure 5F). These results indicate that Edf1 is important for RGC commitment and differentiation towards the multiciliated ependymal cell lineage. Given that Edf1 is consistently expressed in the nucleus from RGCs to MCECs (Figure 5-figure supplement 1A), to further explore how Edf1 regulates the transcriptional level of ependymal multiciliation development process, we used RNA-seq to analyze the change in of ependymal cell development. On the sixth day of differentiation, there were 168 genes up-regulated and 277 genes down-regulated in *Edf1* knockout ependymal cells compared with wild-type. (Figure 5-figure supplement 1B and Figure 5-figure supplement Table 1) We observed that the down-regulated genes were related to the ciliogenic program, such as cilia movement, cilia assembly, and multiciliated cell differentiation (Figure 5D). More notably, among these down-regulated genes, early-phase ciliary transcriptional network-related genes were significantly down-regulated(Choksi et al., 2014; Lewis & Stracker, 2021) (Figure 5E). Consistent with the previous observation of impaired cilia beating and disruption of CSF direction, many key ciliary genes that were critical for cilium movement or planar cell polarity were also down-regulated in *Edf1* knockout ependymal cells (Lewis & Stracker, 2021; Muniz-Talavera & Schmidt, 2017) (Figure 5-figure supplement 1F).

**Figure 5.**
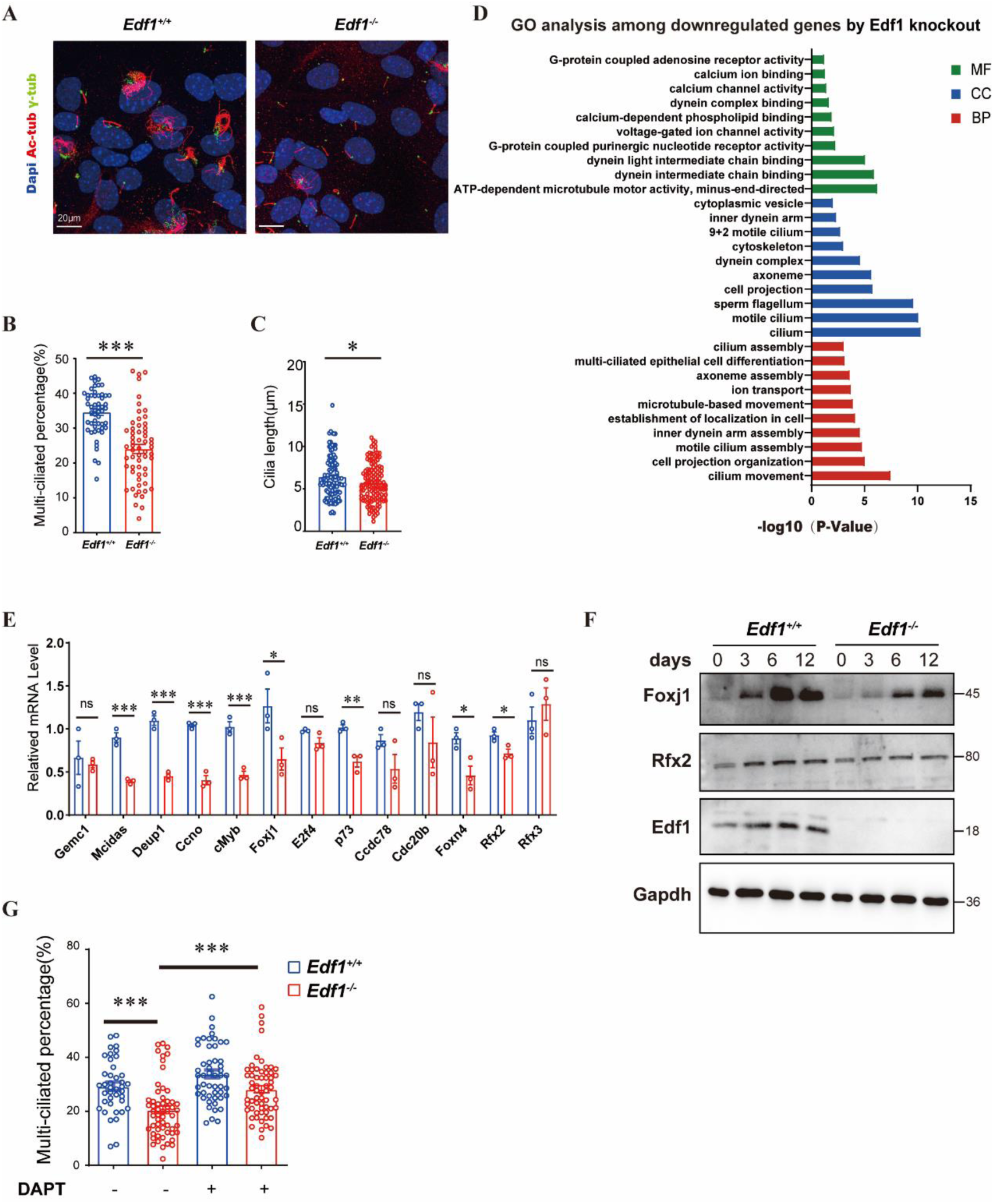
Edf1 controls the MCEC transcriptional networks. (A) RGCs isolated from wild type and *Edf1*^−/−^ P0 mice were cultured in vitro. After 12 days of differentiation, MCECs were immunostaining for Ac-tub(red), γ-tubulin(green) and Dapi(blue). Scale bar: 20 μm. (B-C) The percentage of MCCs that are ciliated based on Ac-tub staining was significantly lower in *Edf1* knockout (n=59) than in wild type (n=53). Cilia length of *Edf1*^−/−^ MCECs (n=128) was shorter than wild type (n=107). Error bars indicate ± SEM. All *P*-values were calculated using an unpaired, two-tailed Student’s t-test, \**P* < 0.05, \*\*\**P* < 0.001. (D) Gene ontology analysis showing top terms significantly enriched among downregulated genes in *Edf1*^−/−^ MCECs after 6 days differentiation. BP: biological process MF: molecular function CC: cellular component. (E) qPCR validation of genes of MCEC transcriptional networks. Error bars indicate ± SEM. n=3, \**P* < 0.05, \*\**P* < 0.01, \*\*\**P* < 0.001, ns. stands for not significant. (F) Detection of Foxj1 and Rfx2 in developing MCECs of wild type and *Edf1*^−/−^ mice by Western blot analysis. (G) The percentage of MCECs of wild type (control n=40; DATP n=50) and *Edf1* knockout (control n=57; DATP n=60) after 12 days of differentiation with or without DAPT treatment. Error bars indicate ± SEM. \*\*\**P* < 0.001

From studies of diverse organisms, inhibition of the Notch signal has become a consistent early event in MCEC differentiation(Lewis & Stracker, 2021). The Notch pathway reduces the ability of ciliary transcriptional networks to promote RGC commitment to the ependymal cell lineage (Mahjoub, Nanjundappa, & Harvey, 2022). To explore whether the differentiation defect of MCECs induced by *Edf1* knockout depends on the downregulation of ciliary transcriptional network-related genes, we used DAPT, which can induce inhibitory effects on Notch 1 signal transduction to prevent the cleavage formation of Notch1 intracellular domain (NICD) during MCEC differentiation (Guseh et al., 2009; Tsao et al., 2009). After DAPT treatment, there is a certain degree of recovery in the proportion of multiciliated cells decreased due to *Edf1* knockout (Figure 5G, Figure 5-figure supplement 1C). This indicates that inhibiting the Notch1 signaling pathway can partially rescue the developmental defects of MCECs caused by *Edf1* knockout. It is worth noting that DAPT treatment can enhance the development of MCECs but does not affect Edf1 expression during development, which indicates that Edf1 expression is not regulated by the Notch1 signaling pathway (Figure 5-figure supplement 1D). In addition, Edf1 knockout did not affect the Notch1 signaling pathway (Figure 5-figure supplement 1D and G). These results suggest that Edf1 may function as one of the most upstream activators of early-phase ciliary transcriptional networks besides the Notch1 signal and maintaining ciliary gene expression at a later phase.

## Discussion

The proliferation and differentiation of neural stem cells are spatiotemporally regulated during brain development. Defects in the development or differentiation of neural stem cells at different stages can lead to congenital hydrocephalus, especially in the development process of MCECs (Ji et al., 2022; Wallmeier et al., 2020). In the present study, we found that Edf1 promotes the differentiation of MCEC by positively regulating the genes of the transcriptional regulatory network related to the development of MCECs. Our results reveal the important function of Edf1 in brain development.

The differentiation of MCCs relies on a unique transcriptional regulatory network. Multiciliation as a conserved process is found throughout evolution. It is still unclear whether conserved mechanisms exist responsible for the multiciliation in both unicellular and multicellular organisms. However, most genes that have been studied in the ciliary transcriptional networks like *Mcidas* are limited to Vertebrata, and the others only present orthologs in Metazoa (Defosset et al., 2021). Given these genes are absent in unicellular organisms, it seems unlikely. Edf1 is highly conserved in diverse organisms including various unicellular species such as *Tetrahymena* and *Chlamydomonas,* which offers the possibility that Edf1 function as an ancient transcriptional co-factor modulates multiciliation in both unicellular and multicellular organisms.

Edf1, as a transcriptional co-factor, participates in the differentiation process of various cells (Leidi et al., 2009; Mariotti et al., 2000; K. Takemaru et al., 1998; Ki Takemaru et al., 1997). Unlike the decrease in Edf1 protein levels during endothelial cell differentiation, from RGC to mature MCECs, Edf1 content does not change significantly in the nucleus. Given that Edf1 activity in endothelial cells is active upon activation of protein kinases C and A (Ballabio et al., 2004; Mariotti et al., 2000), it will be interesting to identify whether Edf1 is regulated by post-translational modifications in the regulation of MCECs development. Considering that Edf1 lacks a DNA binding domain (Ki Takemaru et al., 1997), it is still worth further studying which transcription factors interact with Edf1 to regulate the development process of MCECs.

Although *Edf1^-/-^* mice exhibit a high probability of postnatal hydrocephalus and runted, which exhibits a phenotype similar to ciliary transcriptional relative genes knockout mice (Boon et al., 2014; Lemeille et al., 2020; Terré et al., 2016), the differentiation defect of ependymal multicilia cells caused by *Edf1* knockout is more inclined to early differentiation delay rather than complete absence of cilia. This is consistent with our observation that Edf1 knockout cannot completely suppress the expression of genes such as *Mcidas*. However, unlike Mcidas, which is mainly expressed in the early development of ependymal cells from E14-P7 as early transcriptional activators (Kyrousi et al., 2015), Edf1 continues to be expressed from RGCs to MCECs. *Edf1* knockout leads to the downregulation of cilia swing-related genes and planar polarity-related genes (Ha, Lindsay, Timms, & Beier, 2016; Muniz-Talavera & Schmidt, 2017) with physiological defects of MCECs in vivo. These results suggest that Edf1 not only participates in the activation of the transcriptional regulatory network required for early ependymal cell differentiation but also regulates the expression of genes related to the physiological functions of multiciliated cells.

In conclusion, our study demonstrates that Edf1 controls the differentiation and function of MCECs through activating the transcriptional regulatory networks. Most importantly, the mouse model reveals a previously unknown function of Edf1 in the development of multiciliated ependymal cells, which is crucial for brain development in vivo. Our data make Edf1 an ideal ancient transcriptional activator to participate in the development process of multiciliated cells. In addition, it may help identify human gene mutations related to diseases associated with multiciliated cell development.

## Materials and Methods

### Mice

Edf1 knockout mice were generated on the C57BL/6J background using the CRISPR/Cas9 system. Four sgRNAs were designed to target exons 2-4 of the Edf1 gene. The sequences for these are as follows: gRNA1: CTTTAGTTCAGACACATACC, gRNA2: CCATGTGTTTACCAAAAACA, gRNA3: GTAAGCACAAGCCCCCCCTA, and gRNA4: GTATGGCCTCAAGAGGACGA. The Edf1 knockout colony was bred through heterozygous mating. Genotyping was verified through PCR following genomic extraction from mouse tails. Experimental protocols were approved by the ethics committees of Jianghan University. All animals were maintained in a pathogen-free environment with a room temperature of 23 ℃, humidity between 30-70%, and a 12-hour day/night cycle. Mice had free access to food and water.

### Alignment and phylogenetic analysis

The protein sequences were alignment with Clustal X software. The Genbank accession numbers: *Hs*Edf1 (NP_003783.1), *Mm*Edf1 (NP_067494.1), *Xl*Edf1 (NP_001089760.1), *Ce*Edf1 (NP_502166.1), *Cr*Edf1 (XP_001699725.1), *Dr*Edf1 (NP_957039.1), *Dm*Edf1 (NP_001246795.1), *Tt*Edf1 (XP_001011029.1). An evolutionary relationship was inferred using the Neighbor-Joining method with 1000 replicate bootstrap trees. Evolutionary distances were computed with the p-distance method and are in the units of the number of amino acid substitutions per site.

### Reverse transcription-quantitative PCR

**(RT-qPCR)** Total RNA was isolated from in vitro induced MCECs of both wild type and Edf1 knockout at indicated time points using TRIzol Reagent (Invitrogen). 1ug of total RNA was reverse transcribed to form cDNA using the HiScript II Reverse Transcriptase kit (Vazyme, Cat.#R223-01) for qPCR. The ChamQ SYBR qPCR Master Mix (Vazyme, Cat.#Q311-02) was used for qPCR. The list of qPCR primers used can be found in Supplement Table 1. Gapdh was used as the reference gene.

### Western blot analysis

Induced MCCs were lysed in RIPA buffer (Beyotime, P0013B) with 1 mM PMSF and 1 mM sodium fluoride (NaF). Nuclear/cytoplasmic protein components were extracted using a nuclear protein extraction kit (Solarbio, Cat.#R0050), and then protein lysates were used for standard immunoblotting analysis. The list of antibodies used as follow: anti-Edf1 (1:500, proteintech, 12419-1-AP), anti-Foxj1 (1:500, D360355, Sangon Biotech), anti-RFX2 (1:1000, K110716P, Solarbio), anti-Gapdh (1:1000, ABclonal, AC002) and anti-Notch1 (1:500, Servicebio, GB111690) anti-Histone H3 (1:1000, ABclonal, A2348).

### Histological processing and Immunofluorescence

Mice under the indicated conditions were anesthetized and then perfused with PBS containing 10ug/ml heparin. The brain was dissected out and fixed in 4% PFA at 4℃ overnight. The brains were washed three times for 30 minutes each with PBS, then incubated in 30% sucrose overnight. After dehydration treatment, the brains were processed for cryo-sectioning. A 10um coronal section in the brains of wild type or Edf1 knockout mice at indicated ages was collected at 100um intervals from the appearance of the lateral ventricle to the end of the fourth ventricle. The sections were used for both H&E staining and Immunofluorescence.

For H&E staining, sections were performed by standard protocols. Images were taken at Digital Pathology System (Thermo&3DHISTECH, MIDI) with a 20x objective lens. Images were processed through the software CaseViewer 2.4.

For Immunofluorescence, the cryosections were washed with PBS with 0.2% Tween-20 (PBST)(Sigma) at room temperature for 5 minutes, then permeabilized in a slide-holder containing PBS with 0.5% Tween-20(Sigma) for 30 minutes, and then each slide was incubated with PBST containing 0.3M glycine (Sigma) at room temperature for 30 minutes. Next, blocking was performed with PBST with 5% heat-inactivated donkey serum (sigma). Primary antibodies were incubated overnight at 4℃ then secondary antibodies were incubated at room temperature for 2 hours. Confocal images were taken on a Leica SP8 with a 63x oil immersion objective lens (NA 1.40). Primary antibodies were used as follow: anti-Edf1 (1:500,proteintech 12419-1-AP), anti-Foxj1 (1:500, Sangon Biotech D360355), anti-Pax-6 (1:500, Samta cruz sc-81649), anti-NeuN (1:500, proteintech 66836-1-IG), anti-AQP1 (1:300; ABclonal A15030), anti-γ-Tubulin (1:1000, Merck T6557), anti-β-Catenin (1:300, ABclonal A11932),anti-Acetylated Tubulin, (1:1000, Sigma-Aldrich T6793), anti-S100-B, (1:500, Sigma-Aldrich S2532), anti-ki-67, (1:1000, invitroGen 14-5698-80). Secondary antibodies were conjugated to Alexa Fluor dyes (donkey or goat polyclonal, 1:500, Invitrogen).

### Hydrocephalus incidence

Hydrocephalus was assessed through direct brain anatomical observation. The incidence was quantified by comparing Edf1 knockout mice with their age-matched littermate wild-type controls.

### SEM

The lateral ventricles of wild type and Edf1 knockout littermates at P3 and P5 were dissected out and fixed in Kamovsky’s fixative (2.5% glutaraldehyde, 2% PFA, 0.1M cacodylate buffer, pH 7.4) at 4℃ for more than 2 days. The tissues were then washed with 0.1M Mcacodylate buffer containing 2.5% glutaraldehyde and 1 mM calcium chloride three times for 10 minutes each. Next, the tissues underwent gradient dehydration with ethanol, followed by two washes with HMDS for 5 minutes each. After the preparations were dried, the samples were coated with gold/palladium. Final images were taken on an ultra-high resolution cold field scanning electron microscope (Hitachi, SU8010).

### In vitro ciliogenesis induction and ciliogenesis quantification

The induction of ciliogenesis in MCCs has been previously described (Delgehyr et al., 2015). The brains of P0-P3 wild type or Edf1 knockout mice were dissected in cold-filtered Hank’s Buffer (161mM NaCl, 5mM KCl, 1mM MgSO4, 3.7 mM CaCl2, 5 mM Hepes, and 5.5 mM Glucose, pH 7.4). The olfactory bulbs were removed, and the dissected telencephalon was cut into pieces. Each brain was then added to 1ml of digestion solution (10U/ml Papain, 0.2 mg/ml L-Cysteine, 0.5 mM EDTA, 1 mM CaCl2, 1.5 mM NaOH, and 0.15% DNase I) and incubated at 37℃ for 45 minutes. The solution was centrifuged at room temperature at 1400rpm to collect the precipitate, which was then resuspended in DMEM medium containing 10% FBS to terminate digestion.

Next, the cells were resuspended in DMEM containing 10% FBS after centrifugation and then spread onto a 25-cm2 flask pre-coated with Poly-L-Lysine. Weakly attached cells were removed the following day. After 4-5 days, the cells reached confluence. The cells were digested with trypsin-EDTA solution and re-plated onto a confocal dish pre-coated with Poly-L-Lysine and cultured overnight.

The following day (day 0), the culture medium was replaced with DMEM/Glutamax, 1% P/S without FBS to start induction. Cells were collected at indicated time points thereafter.

For ciliogenesis quantification, MCECs of wild type or Edf1 knockout that had been induced for 12 days were fixed and then stained for Ac-tub and Dapi. Images were taken on a Zeiss Axio Vert A1 microscope with a 100x oil immersion objective lens (NA 1.40). Ciliated percentage was calculated with ImageJ. For the measurement of cilia length, the cilia were imaged as Z-stacks from the base to the ciliated apex at an interval of 0.3um using a Leica SP8 with a 63x oil immersion objective lens (NA 1.40) and a 2.0 zoom-in. The images were then reconstructed in 3D using Imaris software 9.9. The length of the cilia was measured using the filament measurement tool of Imaris software 9.9. Each length value represents the average length of 3 to 4 cilia measured on one MCEC.

### Cilia beats and flow quantification in the brain section

The measurement of cilia beats was described elsewhere (Al Omran, Saternos, Liu, Nauli, & AbouAlaiwi, 2015). For high-speed imaging of ciliary beating, fresh coronal brain sections of P5 wild type or Edf1 knockout mice were placed in 30 mm glass-bottom culture dishes containing DMEM/High-Glucose supplemented with 10% fetal bovine serum (FBS) and 1% penicillin/streptomycin solution (P/S). Live imaging was recorded with a 10 ms exposure time at 100 fps for 3 sec at room temperature using a Nikon ECLIPSE Ti2 microscope, APO 100x oil-immersion lens (NA 1.49), sCMOS high-speed camera, and NIS-Elements software. The frequency of ciliary beating was determined by decreasing the speed of videos and counting the number of beats per second. Each beat frequency represents the average rate of ciliary beating in a given field of view. The beat display diagram shows the delayed image of a cluster of cilia beating within one second. Movies of beats were generated with a frame rate of 25 fps. For the ependymal flow assay (Mirzadeh, Doetsch, Sawamoto, Wichterle, & Alvarez-Buylla, 2010), freshly collected coronal brain sections of P15 wild type or Edf1 knockout mice were placed in a confocal dish containing preheated (37°C) DMEM/High-Glucose supplemented with 10% FBS and 1% P/S solution. Fluorescent microspheres with a diameter of 0.5µm, which had been diluted, were added to the LV region. The movement of the fluorescent microspheres was recorded with a 30 ms exposure time at 100 fps for 30 seconds using a Nikon ECLIPSE Ti2 microscope, Nikon Plan Fluor 20x N2 lens (NA 0.5), sCMOS high-speed camera, and NIS-Elements software. The speed and path of the microspheres’ movement were calculated using Imaris software 9.9. Movies of the fluorescent microspheres were generated with a frame rate of 25 fps.

### RNA-seq and analysis

The RNA-seq was carried out by Wuhan IGENEBOOK Biotechnology Co., Ltd. MCCs of wild type or Edf1 knockout, after 6 days of induction, were collected using TRIzol reagent. Total RNA was extracted with the RNAprep Pure Kit DP432 (TIANGEN Biotech Co., Ltd., Beijing, China), following the manufacturer’s instructions. The integrity of all RNA samples was evaluated using the Qsep1 instrument. 3 μg of total RNA was used to construct RNA libraries with the MGIEasy mRNA Library Prep Kit. This process involved polyA-selected RNA extraction, RNA fragmentation, random hexamer primed reverse transcription, and 100nt paired-end sequencing by DNBSEQ- T7. The adapter and low-quality reads were filtered out through cutadapt (version 1.11). Clean reads were mapped to the Mus musculus reference genome (GRCm38/mm10) by Hisat2 (version 2.1.0), allowing up to two mismatches(Su et al., 2019). These genes were aligned against public protein databases; NR (RefSeq non-redundant proteins). We used Featurecount (v1.6.0) for transcript abundance estimation and normalization of expression values as FPKM (Fragments per kilobase of transcript per million fragments mapped), identifying significantly differentially expressed genes with a fold change >2. We performed Gene Ontology analysis using the DAVID database.

### Whole mount staining of and PCP quantification

The whole mount staining procedure for the lateral ventricle wall refers to previously published literature(Mirzadeh et al., 2010). The dissected lateral ventricle wall was fixed overnight at 4℃ in PBST containing 4% PFA. After blocking at room temperature, the tissue was incubated in primary antibodies at 4℃ for 24 hours, washed three times for one hour with PBST, incubated overnight in secondary antibodies, and then washed three times for two hours. Confocal images were taken on Leica SP8 using a 63x oil immersion objective lens (NA 1.40) with 2.0 zoom-in. BB patch angle and BB patch displacement were quantified by ImageJ. BB patch angles were normalized to the average of these angles at 360° in each MCC.

### Statistics and reproducibility

Histological analysis of internal tissues and physiological function tests were conducted using a minimum of age-matched littermate mice. The number of experimental animals was no less than three per genotype at each time point. In vitro experiments were biologically replicated at least thrice, with three experimental repetitions. All results depicted in the bar graphs are articulated as means ± SEM. The means of two experimental groups were juxtaposed using an unpaired two-tailed Student’s t-test via GraphPad Prism. The distributions of vectorial angles were delineated by Origin software. Differences were deemed significant at p < 0.05.

### Data availability

All relevant data for RNA sequencing have been deposited in the Gene Expression Omnibus database under the accession code GSE244940. The authors declare that the data supporting the findings of this study are available within the article and its supplementary materials or from the corresponding authors upon reasonable request.

## Author contributions

H.J. and Z.H. designed experiments; H.J., M.H., M.D., J.H., S.Z., H.W., W.W., C.X., J.X., H.L. and X.Z. performed experiments; H.J., M.H., M.D., J.H. and Z.H. analyzed data; H.J. and Z.H. supervised the project and wrote the manuscript.

## Acknowledgments

This work was funded by the National Key R&D Program of China (grant number: 2020YFA0907400), the National Natural Science Foundation of China (grant number: 32170702 and 82000828), The Research Fund of Jianghan University (grant number: 2023KJZX24), the Wuhan Science and Technology Bureau of Hubei Province of China (20222508770001) and the Major Special Funding Program for First-class Discipline Construction of Jianghan University (grant number: 2023XKZ021).

## Competing interests

The authors declare no competing interests.

## Additional files

**Figure 1-figure supplement Table. 1**

Information on mice of Survival numbers of embryonic and postnatal wild type and *Edf1* knockout mice.

**Supplement Table. 1**

The lists of primers used in the Genome PCR and real-time PCR.

**Figure 5-figure supplement Table. 1**

The lists of differential expression genes of MCECs between wild type and *Edf1* knockout.

**Figure 4-figure supplement Video. 1**

Representative video of the ciliary beating of wild type MCECs in vivo at P5.

**Figure 4-figure supplement Video. 2**

Representative video of the ciliary beating of *Edf1* knockout MCECs in vivo at P5.

**Figure 4-figure supplement Video. 3**

Representative video of the CSF of wild type brain preparations in vivo at P15.

**Figure 4-figure supplement Video. 4**

Representative video of the CSF of *Edf1* knockout brain preparations in vivo at P15.

## Supplementary figures

**Figure 1-figure supplement 1.**
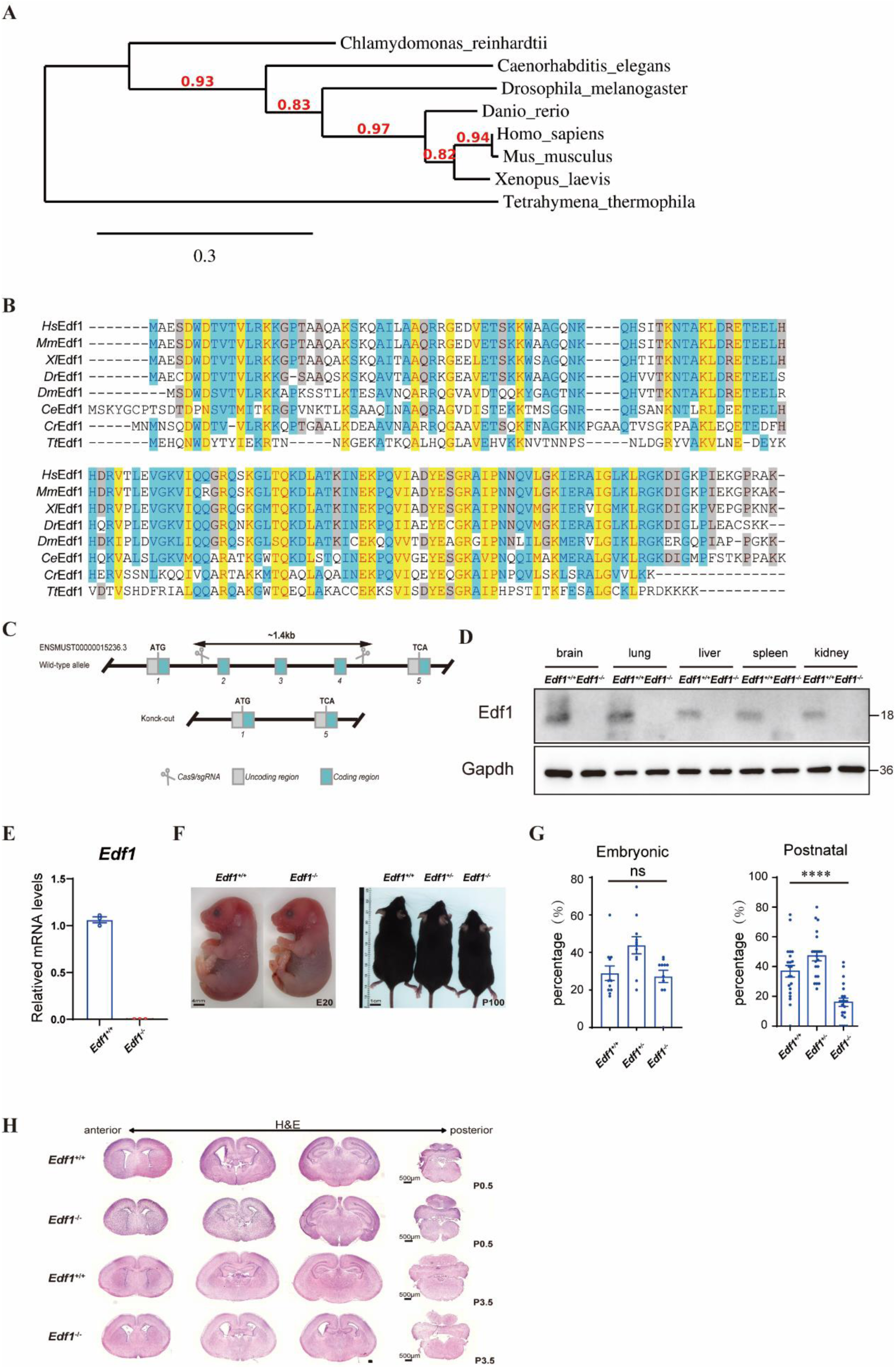
Generation of *Edf1*^−/−^ mice, survival quantities, histological analysis of *Edf1*^−/−^ mice brains at P0.5 and P3.5. (A) Neighbor-joining phylogenetic tree including 8 Edf1 domain-containing proteins from diverse taxa. The tree is drawn to scale, the scale corresponding to the number of amino acid substitutions per site. Bootstrap values are shown near each branch. (B) Alignment of Edf1 domain-containing proteins from various species. (C) *Edf1* gene targeting strategy. (D) Deletion of Edf1 was confirmed at the protein level via immunoblotting in several organs, E. and at the mRNA level of the brain via RT-qPCR. n=3; data are presented as mean values ±SEM. (F) Littermate males of the wild type, *Edf1*^+/−^and *Edf1*^−/−^ at indicated ages. (G) Survival percentages of indicated genotype of embryonic and postnatal animals. Genotyping was taken before birth (n=11) or after P14 (n=21). Error bars indicate ± SEM. All *P*-values were calculated using an unpaired, two-tailed Student’s t-test, \*\*\*\**P* < 0.0001. (H) H&E staining of coronal sections of wild type and *Edf1*^−/−^ mice brains at P0.5 and p3.5. Scale bar: 500 μm.

**Figure 2-figure supplement 1.**
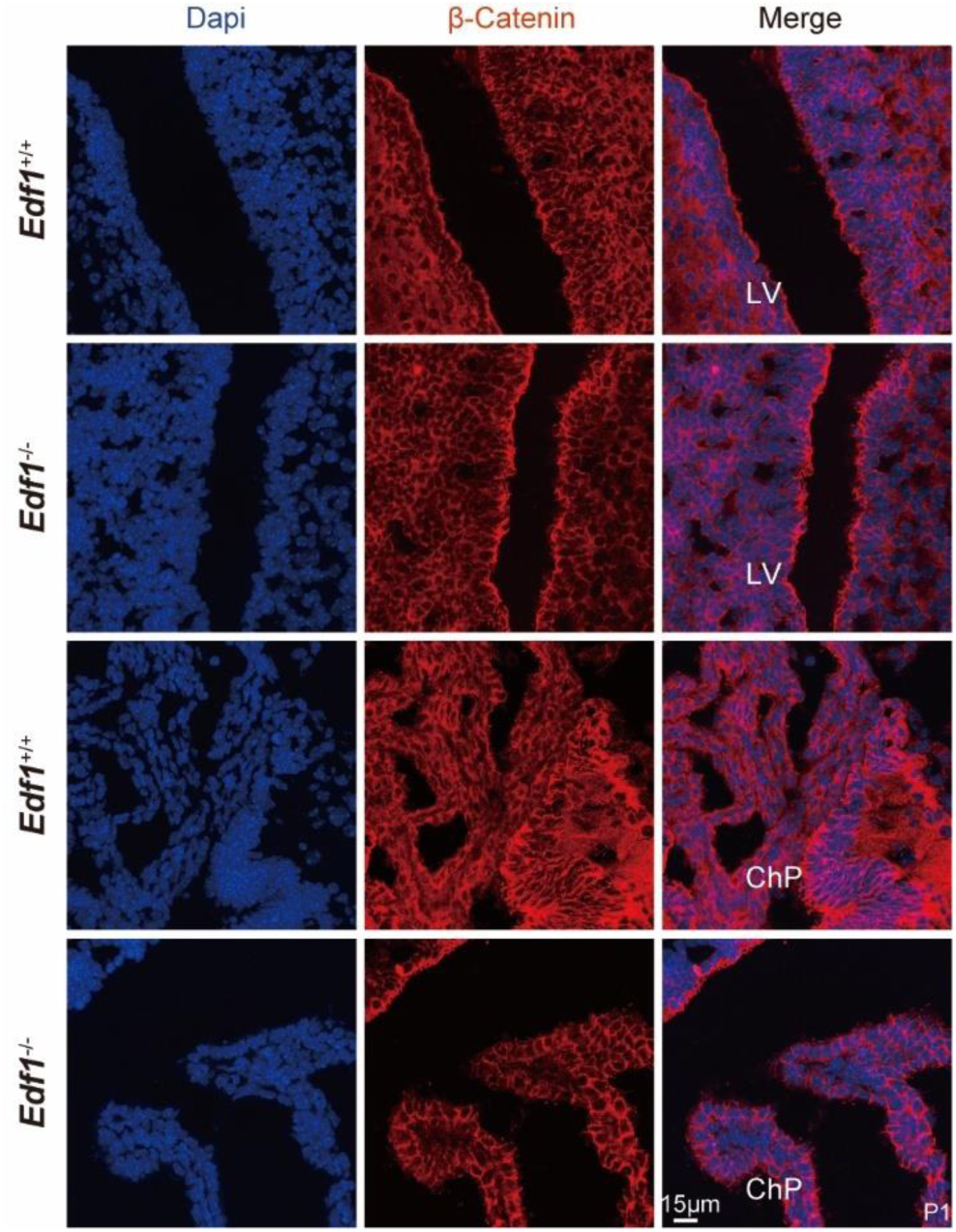
Cell-cell adhesion of LV and choroid plexus. Coronal brain sections from wild type and *Edf1*^−/−^ mice at P1 were stained for anti-β- catenin(red) and Dapi (blue). cell-cell adhesion was not affected. LV lateral ventricle, ChP choroid plexus, Scale bar: 15 μm.

**Figure 3-figure supplement 1.**
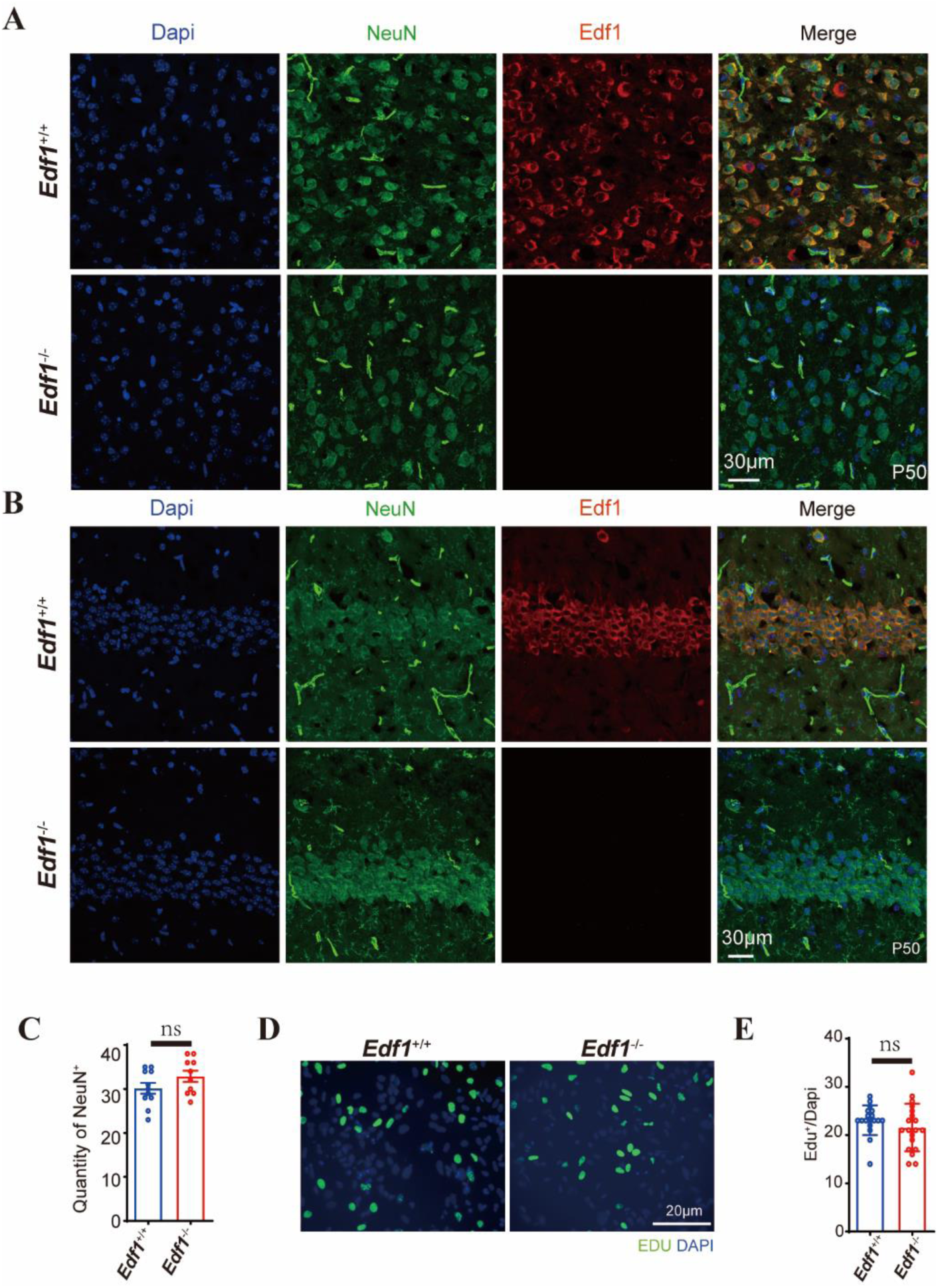
Edf1 co-localizes with NeuN in the cortex and hippocampus. (A-C) Coronal brain sections from wild type and *Edf1*^−/−^ mice at P50 were stained for anti-NeuN(green), anti-Edf1(red), and Dapi (blue). (C) The number of NeuN of CTX(A) and Hip(B) was counted (n = 11 and 10 from 5 wild type and *Edf1^−/−^* mice). Scale bar: 30 μm. (D) and (E) Detection of cell proliferation of wild type and *Edf1*^−/−^ RCGs by EDU immunofluorescence assay (n = 18 and 19 for wild type and *Edf1*^−/−^ mice from 3 independent experiments). Error bars indicate ± SEM. All *P*-values were calculated using an unpaired, two-tailed Student’s t-test. ns., not significant.

**Figure 4-figure supplement 1.**
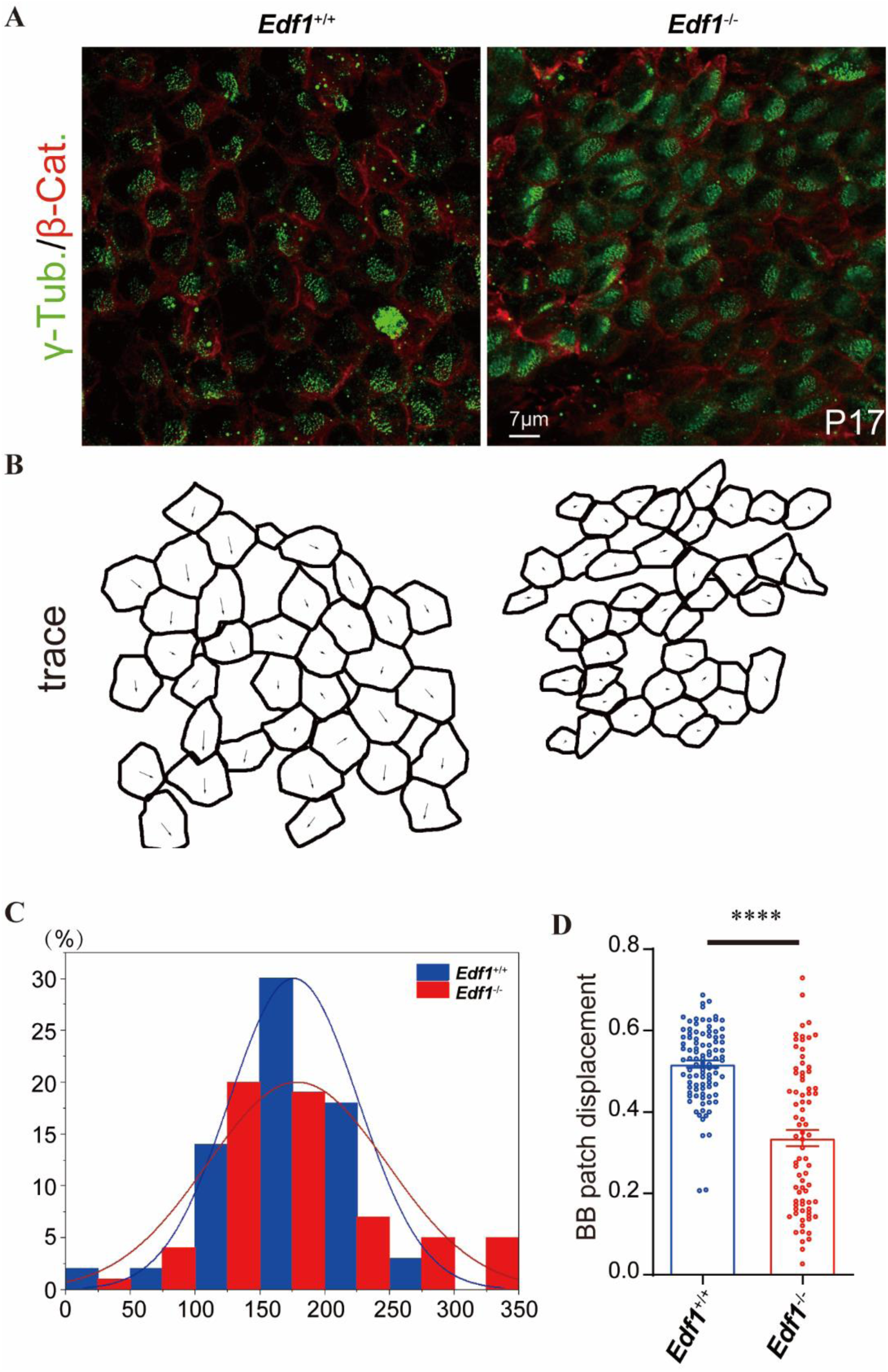
BB patch angle and BB patch displacement were disrupted in *Edf1*^−/−^ mice. (A) Wholemount immunostaining for β-catenin(red) and γ-tubulin (green) of wild type and *Edf1*^−/−^ lateral ventricles at P17. Scale bar: 7 μm. (B) Traces of the intercellular junction labeled with β-catenin of MCCs. The arrows show the angle from the center of the apical surface to the BB patch. (C) The distribution of BB patch angles at P17 was disrupted in *Edf1*^−/−^ mice (n = 61 MCECs from 3 mice) compared with wild type (n = 69 MCECs from 3 mice). (D) Quantification of BB patch displacement at P17 of wild type and *Edf1*^−/−^ mice. BB patch displacement was significantly shorter in *Edf1*^−/−^ mice (n = 78 MCECs from 3 mice) than wild type (n = 97 MCECs from 3 mice). Error bars indicate ± SEM. All *P*-values were calculated using an unpaired, two-tailed Student’s t-test, \*\*\*\**P* < 0.0001.

**Figure 5-figure supplement 1.**
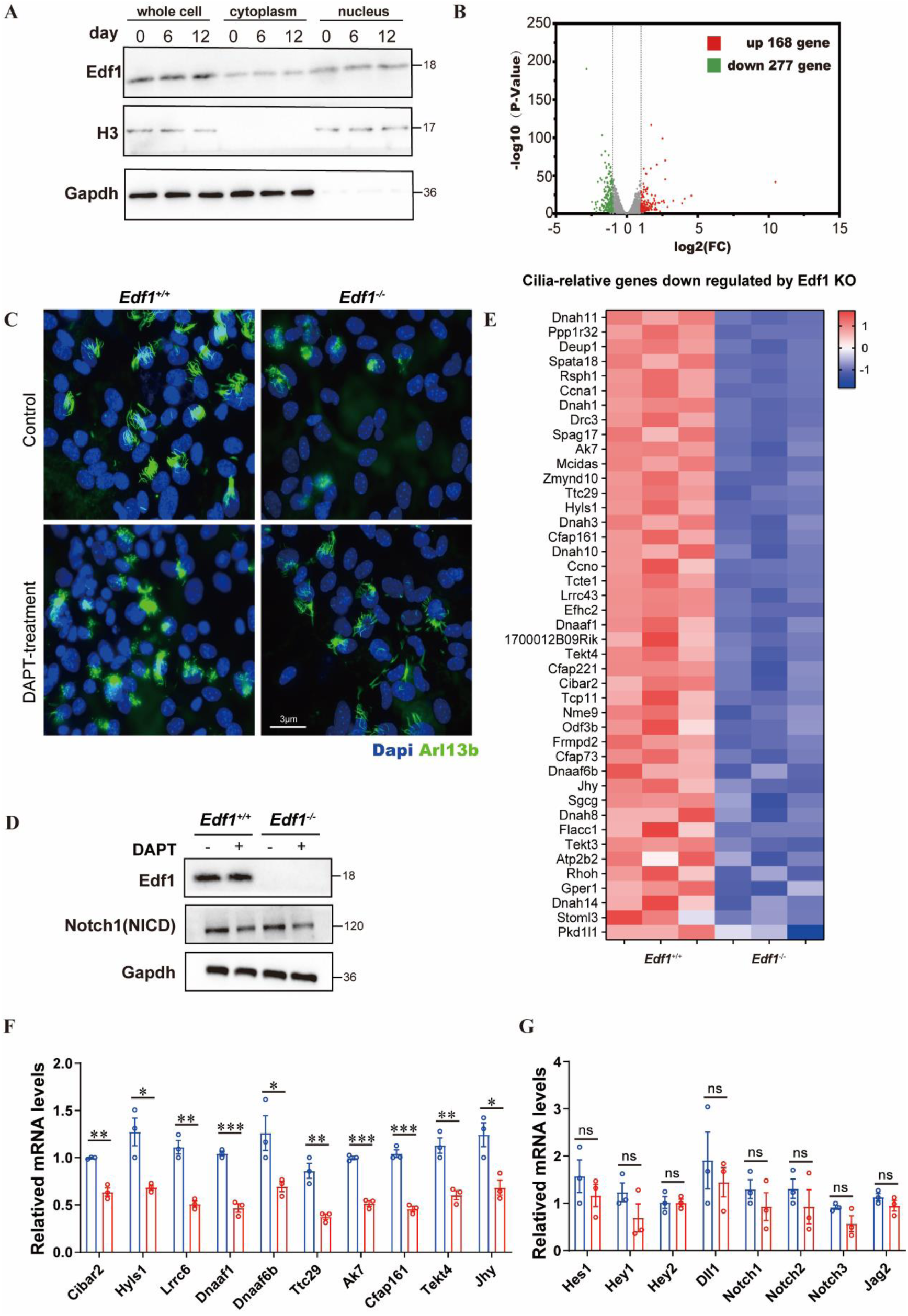
Cilia-relative genes down-regulated by Edf1 knockout. (A) Detection of Edf1 protein from RGCs to MCECs by Western blot analysis. (B) MA plot showing the numbers of differentially expressed genes in Edf1 knockout MCECs. (C) MCECs of wild type and *Edf1* knockout were immunostaining for Arl13b(green) and Dapi (blue) after 12 days of differentiation with or without DAPT treatment. Scale bar: 3 μm. (D) Detection of Edf1 and Notch1(NICD) in MCECs of wild type and *Edf1*^−/−^ mice with or without 10 μM DAPT treatment by Western blot analysis. (E) Heat map shows cilia-relative genes downregulated by Edf1 Knockout. (F) qPCR validation of genes associated with ciliary beating and PCP. Error bars indicate ± SEM. n=3, \**P* < 0.05, \*\**P* < 0.01, \*\*\**P* < 0.001, ns. stands for not significant. (G) qPCR validation of genes associated with Notch signaling. Error bars indicate ± SEM. n=3, ns. stands for not significant.

## Notes

### Competing Interest Statement

The authors have declared no competing interest.

